# What pupil size can and cannot tell about math anxiety

**DOI:** 10.1101/2023.12.15.571819

**Authors:** Elvio Blini, Giovanni Anobile, Roberto Arrighi

## Abstract

**Math Anxiety (MA)** consists of excessive fear and worry about math-related situations. It represents a major barrier to numerical competence and the pursuit of STEM careers. Yet it is still poorly evaluated, mostly through self-reports. Here we sought to probe **Pupil Size (PS)** as a viable biomarker of MA by administering arithmetic problems to young adults (N= 70) with various levels of MA. We found that arithmetic competence and performance are indeed negatively associated with MA, and this is accurately tracked by PS. When performance is accounted for, MA does not further modulate PS (before, during, or after calculation). However, the latency of PS peak dilation could add a significant contribution to predicting MA scores, indicating that high MA may be accompanied by more prolonged cognitive effort. Results show that MA and mathematical competence may be too crystalized in young adults to be discernible, calling for early educational interventions.

## Introduction

Numbers and mathematics are omnipresent in our lives. The ability to grasp mathematical concepts greatly enhances our understanding of an ever complex world, as well as more stable and satisfactory work careers. Conversely, poor numeracy has been associated to early drop-out from school and difficulties in maintaining a stable employment (Bynner & Parsons, 1997). Poor numeracy thus equals fewer opportunities for personal and career growth, with particular groups (e.g., women) that may be disproportionally more vulnerable. Hence, research has increasingly focused on the factors hindering the reaching of an appropriate mathematical competence. The concept of **Math Anxiety (MA)** has gained particular traction. Within the broad domain of anxiety, MA can be defined as excessive fear and worry specific for math-related performance or situations (Caviola et al., 2017; Maldonado Moscoso et al., 2020, 2022). It is a debilitating condition in that it decreases self-confidence in students and creates a barrier that hampers successful learning (Ashcraft, 2002). It is also concerning in that its prevalence has been estimated to be as high as 17% in the general population (Ashcraft & Moore, 2009), or even higher in specific target groups (e.g., students of scientific matters, Betz, 1978); MA may be, however, on the rise following the Covid-19 pandemic and the long phase of remote learning that ensued (Lanius et al., 2022). Phenotyping and tackling MA is thus a key societal priority, as so much is at stake.

One, unresolved problem – at the core of many differences in the reports of MA prevalence and other findings – is that we lack an agreed upon pipeline for optimal identification of MA, including tools and normative criteria. MA is currently only assessed via questionnaires (i.e., the AMAS, Hopko et al., 2003; Primi et al., 2014). Self-reports are handy in that they can be administered in a quick, inexpensive, simple fashion, without the need of particular training, and provide a coarse, but sufficiently reliable assessment of one’s anxiety levels. However, self-reports also have limitations, in that they often require an advanced metacognitive competence and may be subject to biases, e.g. related to social acceptance. For these reasons it would be particularly important to leverage on measures capable to escape awareness, more tightly linked to the physiological processes causing or accompanying MA, and therefore in principle perfectly capable to improve its identification.

Previous fMRI studies have shown that individuals with high MA tend to recruit comparatively more a broad fronto-parietal network devoted to attention and arithmetic performance than individuals with lower MA (Atabek et al., 2022; Chang et al., 2017), even with very simple calculations for which behavioral differences are not apparent (Chang et al., 2017). The reasons why this should be the case are unclear. One possibility is that individuals with high MA may deploy suboptimal strategies for the calculations, less automatic and thus more demanding and reliant on a frontal network for executive functions (Chang et al., 2017). Alternatively, even when deploying the same strategy, high MA may be accompanied by a less efficient information processing, so that the same cognitive operations are prolonged in time and/or more demanding (Chang et al., 2017). This possibility fits nicely with important accounts of MA, such as the cognitive interference theory (Ashcraft & Kirk, 2001), postulating that anxiety effectively adds a continuous dual-task setting that subtracts attentional resources from the calculation, and thus impairs performance. Interference would particularly affect working memory, though emotional components may also concur: people with high MA would additionally need to cope with unpleasant affective reactions to math, further diverting attentional resources away from the main task. Interestingly, (Lyons & Beilock, 2012) reported that increased fronto-parietal activations *preceded* the onset of mental calculation. This finding is difficult to explain in terms of a different numerical competence across different MA levels or different levels of mental effort required. Rather, it may have, to some extent, a proactive nature in that this activation could mitigate subsequent behavioral performance deficits (Lyons & Beilock, 2012). At any rate, studies have generally shown that math-related stimuli quickly and automatically grab attention, well before mental calculation is performed, also through subcortical connections (Lyons & Beilock, 2012; Pizzie & Kraemer, 2017) most notably including the amygdala (Pizzie & Kraemer, 2017). To summarize, despite the limitations due to the small sample size, the physiological manifestations of MA appear rather consistent across studies, and may in principle help increasing the precision in measuring MA. Furthermore, these studies suggest that the calculation stage, while extremely relevant, may not be the only one apt to characterize MA, calling for a more complete and “naturalistic” approach to mental arithmetic.

Most, state of the art physiological recordings (e.g., fMRI, MEG) can be too impractical for many, expensive, and complex to handle. Here, we focus on **pupil size (PS)** which, despite having received comparatively less attention (but see Layzer Yavin et al., 2022; Throndsen et al., 2022), provides a cheap but very informative readout of several cognitive processes (Mathôt, 2018; Strauch et al., 2022), as well as the state of balance of the autonomic nervous system. Measuring PS allows one to assess how mental processes unfold in time, with a reasonable temporal resolution along the course of a single trial. Increased PS is associated with phasic activation of the locus coeruleus (Aston-Jones & Cohen, 2005), a major noradrenergic hub involved in the integration of the attentional networks in the brain, as well as in balancing bottom-up and top-down attention (Reynaud et al., 2021). Far from reflecting passively responses to light (Binda et al., 2013; Binda & Murray, 2015), PS provides insights into automatic orienting processes (Blini & Zorzi, 2023; Castaldi et al., 2021; Salvaggio et al., 2022), emotional states (Bogdanova et al., 2022; Dureux et al., 2021), or working memory load (Ahern & Beatty, 1979; Beatty & Kahneman, 1966; Hess & Polt, 1964; Lisi et al., 2015). Indeed, mental arithmetic has been the task with which the use of PS has been pioneered almost 60 years ago (Ahern & Beatty, 1979; Beatty & Kahneman, 1966), resulting in this measure firmly entering the arsenal of experimental psychologists’ worldwide. It is well established that PS is increased the more arithmetic problems are difficult (i.e., with increased cognitive demands). Furthermore, people with lower scores in scholastic aptitude tests generally present larger increases, indicative of the need to deploy comparatively more effort (Ahern & Beatty, 1979). Finally, classic studies have also described a much smaller, but still noticeable, “alertness” effect preceding the onset of mental calculation (Ahern & Beatty, 1979; Hess & Polt, 1964). This pattern of results is therefore reminiscent of what, with fMRI, has been described as an increased fronto-parietal activation, firmly putting forward PS as a cost-effective alternative measure to study MA. The possible mediator role of MA in modulating PS during mental arithmetic has been, however, seldom studied (but see Throndsen et al., 2022), while evidence of this kind would contribute substantially to our understanding of the processes at play. Here we sought to fill the gap by measuring PS in a large sample of university students (N= 70) with various degree of MA. We had three experimental questions. First, we wondered if MA could modulate PS beyond the impact of mathematical performance. This possibility has been mostly neglected in pupillometry studies, but would provide essential context to what is a seminal finding in the cognitive neurosciences (Ahern & Beatty, 1979). Second, we sought to assess whether PS could explain part of the variance in the scores obtained in the elective questionnaire to measure MA, and to what extent. Positive results would enrich our capability to evaluate MA with a tool complementary to questionnaires. Lastly, we wondered which moment of mental calculation, if any, is the most vulnerable to the impact of MA. As outlined above, the phase *anticipating* mental calculation may represent a stage in which MA is already fully deployed. Anticipation is not, however, the only possible signature of MA. In principle, MA may very well unfold when awaiting a feedback, or upon its reception, moments that are likely very salient when considering that MA may include fears about others’ judgments. For all these reasons we move beyond the state of the art in behavioral approaches by putting forward a paradigm in which all these phases (i.e., anticipation, calculation, expectancy of the feedback, and the feedback itself) are combined.

## Methods

All materials, raw data, and analysis scripts for this study are available through the Open Science Framework website: https://osf.io/szb24/

The core functions for preprocessing and analysis are available through GitHub: https://github.com/EBlini/Pupilla

### Participants

In this work, we planned to acquire pupillary responses at different phases of mental arithmetic – that is before the calculation, during the calculation, following the response and awaiting for the feedback, and the feedback phase itself. The main dependent variable was the **Math Anxiety (MA)** score obtained from the most established questionnaire to this aim (AMAS, Primi et al., 2014, see below). We thus powered our study assuming a multivariate linear regression design and a small-to-medium effect size (partial R^2^) of 0.2, which would constitute a sensible bar for discussing pupil size as a useful indicator of MA. The power analysis suggested that a minimum of 70 participants would be needed to achieve 80% statistical power with a 5% error rate. Power curves are depicted in **Figure 1B**.

**Figure 1:**
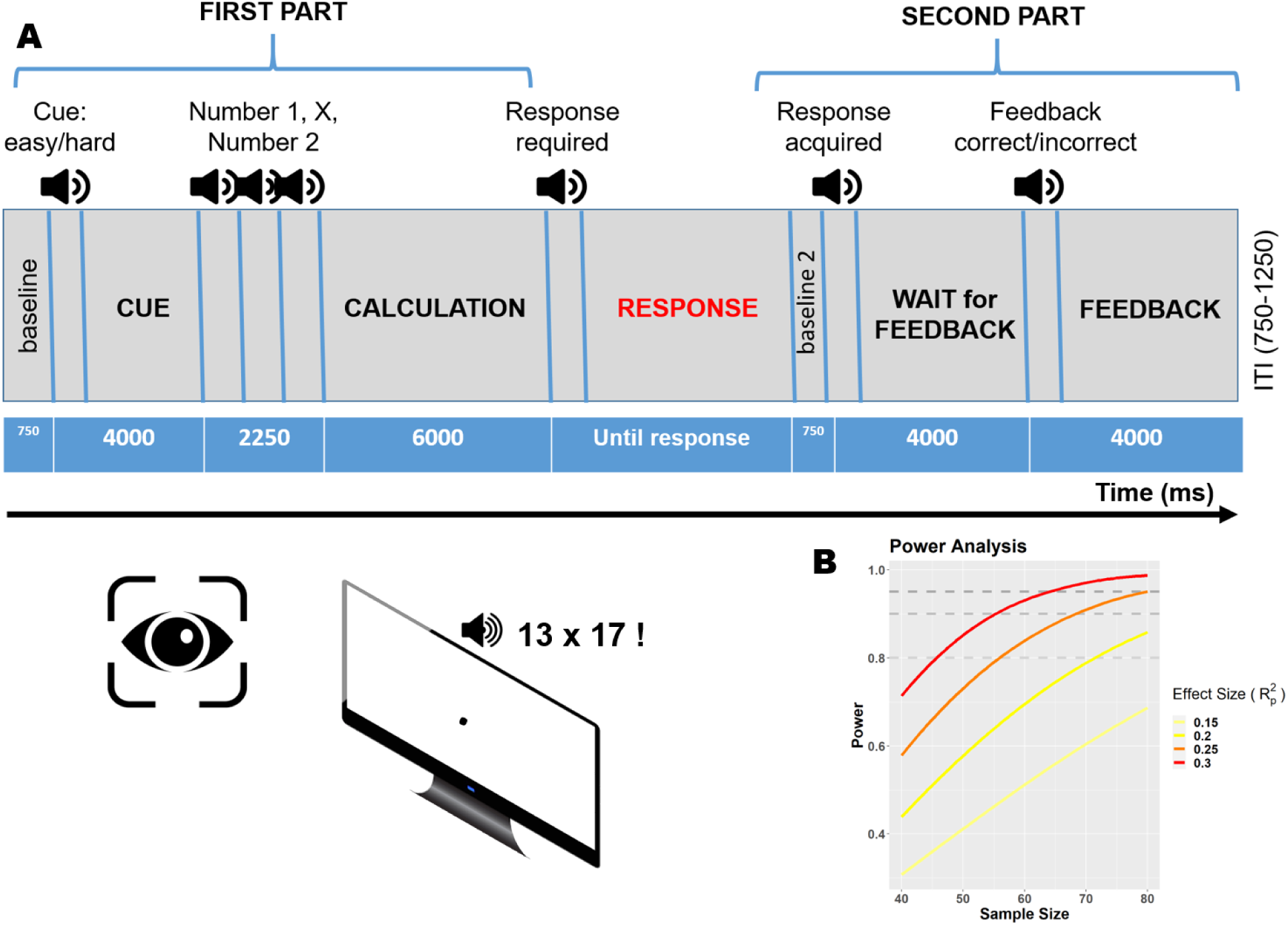
experimental methods. **A)** Graphical depiction and time course of the behavioral paradigm used to assess mental arithmetic. All stimuli were delivered auditorily, hence keeping visual stimulation minimal and constant. The paradigm consisted of several phases: 1) *anticipation*: one word (“easy” or “hard”) is presented auditorily, prompting a problem of corresponding difficulty; 2) *early calculation*: the two numbers and the operator are presented, and mental calculation can start; 3) *response*: a response can be provided verbally in this phase, else the calculation can continue until needed – this makes the traces of variable length after this point; 4) *waiting for the feedback*: a response has been provided, a sound has confirmed that it has been recorded, and the participants are informed that a feedback is upcoming; 5) *feedback*: two different sounds inform the participants about their accuracy. At the beginning of each part, a short (750ms) window was included in order to calculate a baseline for pupil diameter. For all the relevant details and procedures, see the main text. **B)** Power curves as a function of a range of sample and effect sizes.

We therefore enrolled 74 participants overall, all students from the University of Florence. Inclusion criteria were normal (or corrected-to-normal) vision, and no history of neurological, psychiatric, or other sensory disorders. Four participants had to be discarded: 2 for calibration failure; 1 later reported a diagnosis of dyscalculia; 1 did not finish the task and dropped out from the study. The remaining participants were mostly females (N= 49, 70%) with an overall age of M= 23.8 SD= 5.1, range 19-49 years. The experimental procedures were approved by the local ethics committee (Commissione per l’Etica della Ricerca, University of Florence, July 7, 2020, n. 111). The research was carried in accordance with the Declaration of Helsinki, and informed consent was obtained from all participants prior to the experiment.

## Materials and methods

The psychometric assessment battery comprised four established questionnaires. The **Abbreviated Math Anxiety Scale (AMAS)**, Italian version, was administered to quantify MA (Primi et al., 2014). AMAS has been found to be a parsimonious, reliable, and valid measure of MA, invariant across genders, in a sample of high school and college students akin to our pool of participants. The confirmatory factor analysis (Primi et al., 2014) suggested that its latent structure encompasses both *learning MA* – i.e., anxiety related to the process of learning math, such as listening to lectures in class – and *evaluation MA* – i.e., anxiety related to testing situations, such as thinking to an incoming math test. The **Test Anxiety Inventory (TAI)** (Spielberger et al., 1978) was instead administered to quantify anxiety related to general test-taking situations; it requires reporting a range of autonomic symptoms occurring prior to, during or after exams, regardless of their subject. Finally, trait anxiety was measured through the **State-Trait Anxiety Inventory (STAI)** (Pedrabissi & Santinello, 1989). The STAI can be used in a clinical setting to help differentiate anxiety from depression; for this study we only retained the subscale evaluating more stable aspects of proneness to anxiety, i.e. trait anxiety. In addition to MA, test anxiety, and trait anxiety, we sought to operationalize the overall math competences and achievements of our participants through **Mathematical Prerequisites for Psychometrics (PMP**: Prerequisiti di Matematica per la Psicometria; Galli et al., 2011), a paper and pencil test (in Italian) devised to evaluate one’s competence in distinct pillars of mathematical reasoning (e.g., equations, relations, fractions, logic). The PMP was used to probe the impact of MA on lifelong numerical competence alongside in-task measures (i.e., accuracy rate). All these questionnaires were administered prior to exposure to the experimental task.

Once in the lab, participants were tested in a dimly lit, quiet room, their head comfortably resting on a chinrest. They faced a remote infrared-based eye-tracker (EyeLink 1000, SR Research Ltd.), at a distance of approximately 57 cm from the screen. The session started with a 15-points calibration of the eye-tracker, which was then set to monitor participants’ pupil size continuously at a 500 Hz sampling rate. The open-source software OpenSesame (Mathôt et al., 2012) was used to display one fixation dot on screen, present auditory stimuli, and record participants’ responses, along the procedure outlined below and illustrated in **Figure 1A**.

### Procedure

#### Paradigm

Being chiefly interested in pupillary responses, and being pupil size strongly affected by even small changes in environmental light, we opted for a purely auditory set of stimuli. The only visual stimulus on screen throughout the entire testing session was a small gray dot (0.5°) on a black background, which was designed to help and constrain central fixation. The structure and time course of experimental trials is depicted in **Figure 1A**. Each trial started with a 750ms window useful to set a trial-wise baseline pupil size. Then, one of two words (“easy” or “hard”) were acoustically presented, cueing the difficulty of the subsequent mental calculation. This window lasted 4000 ms since the cue onset, and was meant to measure the anticipatory autonomic reactions to the prospect of mental arithmetic. It was followed by 3 sounds, each lasting 750 ms, providing in order the first number (multiplicand), the operation sign (“times”, as only multiplications were provided), and the second number (multiplier). Given these elements, participants were in the position to mentally perform the multiplications. They were asked to do so accurately and while trying to maintain central fixation. Starting from the offset of the second number, there was a 6000 ms time window in which participants were asked *not* to provide an answer. This was to avoid possible artifacts arising from the response, which was provided vocally. Participants were therefore informed about a *minimum* response time limit, which was clearly indicated by the presentation of a “beep” sound, though they were also told that there was no *maximum* time limit for their responses. This feature was introduced to discourage participants switching from precise arithmetic to heuristics, guessing, or estimation strategies, which do not yield similar levels of cognitive effort and thus are not so closely tracked by pupil dilation. Once a response was provided, the experimenter scored it manually in the program.

Thus, this part had a variable duration, depending on the participants’ and experimenters’ response times; furthermore, blinks were allowed in this part of the trial, in consideration of its long total duration, to minimize participants’ discomfort. Participants’ responses end what we refer to “First part” of the trial, in which responses can be aligned precisely with the onset of the trial. In what we refer to as “Second part”, a response has been provided by participants but, due to different response times, the pupillary traces up to that point have different length and duration. The second part begins with a 750 ms sound (a “cash register”), informing participants that their responses had been, indeed, recorded. This phase lasted 4000 ms for the onset of the sound and was meant to measure autonomic pupillary reactions occurring when awaiting for a feedback about one’s own performance, i.e. the anticipatory reactions to the prospect of evaluations. Finally, one of two sounds (each associated to correct or incorrect responses) were delivered for 750 ms while autonomic and affective reactions to the feedback itself were recorded (4000 ms total). Each trial additionally included an inter trial interval randomly set to last between 750 and 1250 ms (uniform jitter). All participants underwent four practice trials, discarded for analyses, prior to the experimental testing, composed of 72 trials administered in two blocks and separated by a pause.

#### Stimuli

We capitalized on the same stimuli used in both classic studies on mental effort and pupil dilation (Ahern & Beatty, 1979) and more recent attempts to study MA (Throndsen et al., 2022). In particular, we selected 12 “easy” and 12 “hard” multiplication problems; note that the contrast “easy vs hard” was the only one reaching significance in the recent study by (Throndsen et al., 2022), suggesting it was the most useful to provide a proxy for cognitive effort. Furthermore, the choice of these two categories allowed for the multiplicands (in addition to the sign operator) to be the same for both categories, thus reducing perceptual (auditory) differences between conditions. Specifically, all operations involved a multiplicand between 12 and 14; for “easy” trials, the multiplier varied between 6 and 9, whereas for “hard” problems the multiplier ranged between 16 and 19. Each combination of these numbers was repeated three times in random order, thereby accounting for 72 overall trials. All auditory stimuli were created offline by means of a vocal synthesizer, edited to last about 750 ms, and then normalized to the same intensity. While auditory traces have been processed to eliminate low-level confounds as much as possible, each stimulus (e.g., numbers, cues) present spectral differences that are unavoidable when conveying different words. Note, however, that the crucial test of this study involves correlations between participants’ reactions and their questionnaires, so that the effect of low-level perceptual differences between sounds on pupil size, if any, is of secondary importance given that the outcome measures were the correlations with MA levels, and all sounds were presented as such to all participants.

### Data preprocessing

Practice trials were discarded. In addition to the blinks automatically identified by the eye-tracker, which were discarded, we further used a velocity-based criterion to identify likely artifacts; each gap in the traces was then extended by 100 ms on each side (before and after the artifacts) before moving to the next steps. Trials in which more than 40% of the data were missing were excluded, and the gaps in the remaining ones were linearly interpolated. Traces were then smoothed through cubic splines (i.e., low-pass filtered). Next, traces were down-sampled to 25 ms epochs by taking the median pupil diameter for each time bin. We calculated the baseline pupil size to use as a covariate in the main analyses in order to probe whether tonic, as opposed to phasic changes in pupil size are related to MA or performance; to this aim, we averaged the pupil size of all participants limited to the first fixation phase, that is a baseline without active tasks or stimuli, measured in arbitrary eye-tracker units. In order to better cope with inter-individual differences we then z-tranformed pupil diameter values separately for each participant (Blini & Zorzi, 2023; Dureux et al., 2021). With normalization, a value of 0 represents the subject-specific mean pupil diameter and, regardless of baseline values, scores represent the relative pupil size expressed as a fraction of the overall participant’s variability. This is useful to account for the fact that the same amount of relative dilation or constriction (e.g., 0.1 mm) has very different meanings in participants with small vs. large baseline pupil size. Finally, all series were realigned to the first baseline window by subtraction of the corresponding, trial-wise pupil size. Trials with extreme baseline values (below -2 or above 2 standard deviations) were discarded. Overall, data cleaning led to discard an average of 5.6% of trials per participant (SD= 4%, range 1.4-22.2%), thus leaving sufficient trials per cell for the main analyses.

In addition to assessing pupil size for each 25 ms timepoint along the course of the entire trial, we obtained summary variables quantifying different aspects of pupil dynamics. Tonic activity was indexed by the average pupil size during the first fixation phase, as outlined above. We then calculated: i) the maximum pupil dilation (relative to the baseline), and ii) the moment in time in which pupil size reached this maximum (i.e., **Peak Latency**); these measures, averaged for each participant, provide indices of the extent and duration, respectively, of the cognitive effort required to solve a task.

### Statistical modelling

We started with computing pairwise correlations for questionnaires and relevant variables; following visual inspection of the data, we opted for Pearson’s correlations because variables appeared normally distributed.

Next, we wondered whether MA, test anxiety, or trait anxiety, separately, could: i) partly explain and modulate PS, ii) could do so even when mental calculation accuracy, which turned out to be an excellent predictor in its own right, is accounted for; and iii) in which experimental phase among those delineated by our experimental paradigm. For the sake of simplicity we followed (Mathôt & Vilotijević, 2022) in using crossvalidated **Linear Mixed Effects Models (LMEM)** through the lme4 package for R (Bates et al., 2015). In this approach all trials from each participant are assigned deterministically to one of 3 folds. Two folds are circularly used as the training set; here, intercept-only LMEMs are performed for each timepoint, and the timepoint having the peak t-value (for each fixed effect or interaction) is then used in the test set to confirm the overall consistency of the target factor across folds. This approach is computationally efficient and very powerful in suggesting the presence of a consistent experimental effect *somewhere* along the time course of the trials. In order to enhance the precision in identifying a temporal cluster for any given effect, we additionally scored a consensus between folds, as timepoints in which all folds presented t-values above |2|; timepoints in which this rather stringent, albeit arbitrary criterion was met were thus identified as a temporal cluster. This approach was repeated, for the sake of the exposition, first without and then with in-task accuracy as further independent variable. Because when accuracy was included as a predictor we did not find substantial modulating effects of MA on PS, we also carried the most liberal approach to the data to ensure that was not an artifact of the robust analyses outlined above. Specifically, we ran intercept-only LMEMs for each time-point without any correction for multiple comparisons.

Finally, we sought to predict the anxiety scores (obtained from the questionnaires) with measures of pupillary dynamics (baseline PS, maximum dilation, and peak latency), mathematical competence (PMP), and performance (accuracy rate). The value and latency of PS peak, in particular, were used as measures of the extent and duration, respectively, of cognitive effort. All variables were summarized down to one value per participant through a grand average. In order to avoid overfitting, we opted for a **Best Subset Regression (BSR)** approach using the **Bayesian Information Criterion (BIC)** as main model performance evaluator. BSR consists in testing all the possible combinations of predictors in different linear models, thus shielding from the shortcomings of stepwise multiple regression (e.g., the order of predictors having an impact on the outcome). At the same time, model complexity and the issue of multiple comparisons is taken care by using the BIC as selection criterion, which penalizes model’s complexity and favor a parsimonious solution with good fit to the data. The difference in BICs between models can be interpreted in a straightforward way in terms of strength of evidence under a uniform, non-informative prior (Raftery, 1995). The best features selected in this step were then probed in follow-up, crossvalidated regressions to assess their predictive power. We adopted a Leave-One-(Subject)-Out (LOO) crossvalidation setup, where just one participant was circularly included in the test set, and computed the resulting coefficient of determination cv-R^2^.

## Results

### Psychometric assessment

**Table 1** reports the descriptive statistics of the collected measures. The correlations between the several tests composing the psychometric assessment battery are depicted in **Figure 2A**.

**Figure 2:**
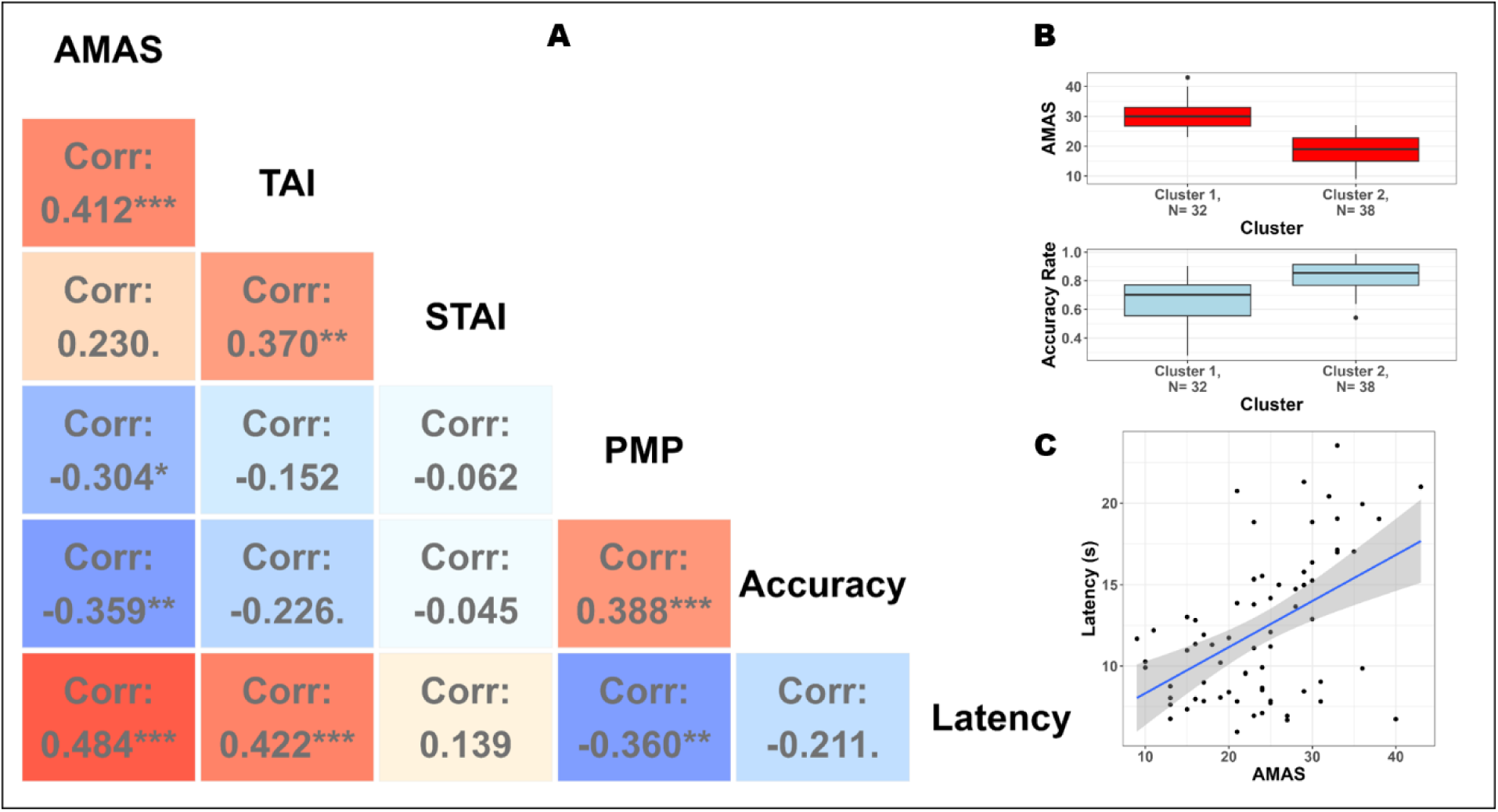
psychometric assessment. **A)** Pearson’s correlations between all the tests included in the assessment and in-task performance measures. Math Anxiety, assessed by the AMAS questionnaire, presents noticeable correlations with other tests evaluating anxiety, in particular Test Anxiety (TAI) but to some extent also Trait Anxiety (STAI). As expected, MA is negatively associated to mathematical competence as assessed by the PMP, confirming that higher MA is associated with poorer numeracy overall. Finally, MA is significantly associated to mathematical performance: higher MA resulted in lower accuracy rate to arithmetic problems. There was a substantial correlation between MA and the latency of pupillary peak dilation, indicating that cognitive effort is prolonged in individuals with high MA (and decreased performance). Symbols: **\*\*\*= p<.001; **= p<.01; *= p<.05; .= p<.1**. **B)** MA and mathematical performance are extremely intertwined. We used unsupervised k-means clustering to show that it is parsimonious to assume that individuals with high MA also have decreased efficiency (task accuracy) on average, with distributions that show little overlap between the two measures. **C)** Details and depiction of the correlation between MA and the latency of the peak pupillary response (time since the offset of the second number, in seconds).

**Table 1.** Descriptive statistics of the collected measures.

| tests | Mean | Median | SD | Min | Max | Skew | Kurtosis |
| --- | --- | --- | --- | --- | --- | --- | --- |
| **AMAS** | 23.84 | 24 | 7.64 | 9 | 43 | 0.13 | -0.52 |
| **TAI** | 44.01 | 45.5 | 12.44 | 21 | 71 | 0.27 | -0.92 |
| **STAI** | 46.06 | 44 | 9.97 | 30 | 66 | 0.3 | -0.98 |
| **PMP** | 24.59 | 26.5 | 5.1 | 11 | 30 | -1.3 | 0.8 |
| **ACC** | 0.75 | 0.79 | 0.16 | 0.28 | 0.99 | -1.01 | 0.38 |
| **Pupil Latency** | 12.25 | 11.33 | 4.47 | 5.97 | 23.56 | 0.61 | -0.69 |
| **Max Dilation** | 2.6 | 2.55 | 0.53 | 1.66 | 4.48 | 0.8 | 1.06 |

Math Anxiety, as assessed by the AMAS questionnaire was moderately and positively related to test anxiety (TAI: r_(68)_= 0.41, *p*< .001, c.i.[0.2 – 0.59]), and less so to trait anxiety (STAI: r_(68)_= 0.23, *p=* .055, c.i.[0 – 0.44]). As expected, there was a negative correlation between MA and mathematical competence (PMP: r_(68)_= -0.3, *p*= .01, c.i.[-0.5 – -0.07]): individuals with higher MA were also those with less developed competences related to math (beyond mental calculation, including logic, relations, etc.). With respect to math performance, people with high MA were also less accurate on average (Accuracy: r_(68)_= -0.36, *p=* .002, c.i.[-0.55 – -0.14]).

To better understand the tight link between MA and performance in terms of accuracy, we used an unsupervised clustering algorithm (k-means) with the objective of assessing whether natural subgroups of individuals can be identified within the entire sample. Clustering has indeed the objective of delineating k groups with the constraint that differences must be maximized between groups and minimized within groups – i.e. groups must be homogeneous internally but also clearly separated. The solution with k= 2 tended to group together participants with high MA and lower task accuracy, which were clustered separately from participants with low MA and higher accuracy. As can be appreciated in **Figure 2B**, there was little overlap between the distributions of the two variables between different clusters. This analysis thus strengthens the notion that MA and arithmetic competence are strictly intertwined, at least in the case of young adults.

We then computed the correlation between MA and the average latency of the peak pupillary dilation. There was a clear positive correlation between MA and PS latency (Latency: r_(68)_= 0.48, *p<* .001, c.i.[0.28 – 0.65], **Figure 2C**): the higher MA, the more sustained and prolonged pupil dilation, with the peak occurring much later in time. Note that a more prolonged pupil dilation can be taken as a proxy of a more prolonged cognitive effort, indicating that students with high levels of MA need to invest in mental calculations longer than peers with lower anxiety scores. The same is true for people with high levels of test anxiety (r_(68)_= 0.42, *p<* .001, c.i.[0.21 – 0.6]) but not necessarily trait anxiety (r_(68)_= 0.14, *p=* .25, c.i.[-0.1 – 0.36]), suggesting that this feature may be shared by the more performative domains of anxiety.

### Does math anxiety modulate pupil size?

First, our stimuli and paradigm were effective in including (and cueing) problems of different difficulty. Hard trials, cued by the respective word, were indeed associated with lower accuracy (accuracy: 61%±26 vs. 90%±9; *t*_(69)_= 10.9, *p*<.001). In terms of pupillary dynamics, hard trials were also associated with a larger maximum pupil dilation (z-scores: 2.83z±0.6 vs. 2.38z±0.53; *t*_(69)_= 8.28, *p*<.001) occurring on average much later in time (seconds: 15.5s±6.4 vs. 8.9s±3; *t*_(69)_= 12.6, *p*<.001). These results thus indicate that the operations that we a priori identified as being easy or difficult were indeed performed as if this was the case.

The time course of pupil dilation is depicted in **Figure 3A**. The plots separate individuals with high or low MA by means of median split, but all the relevant analyses were carried out in a parametric fashion.

**Figure 3:**
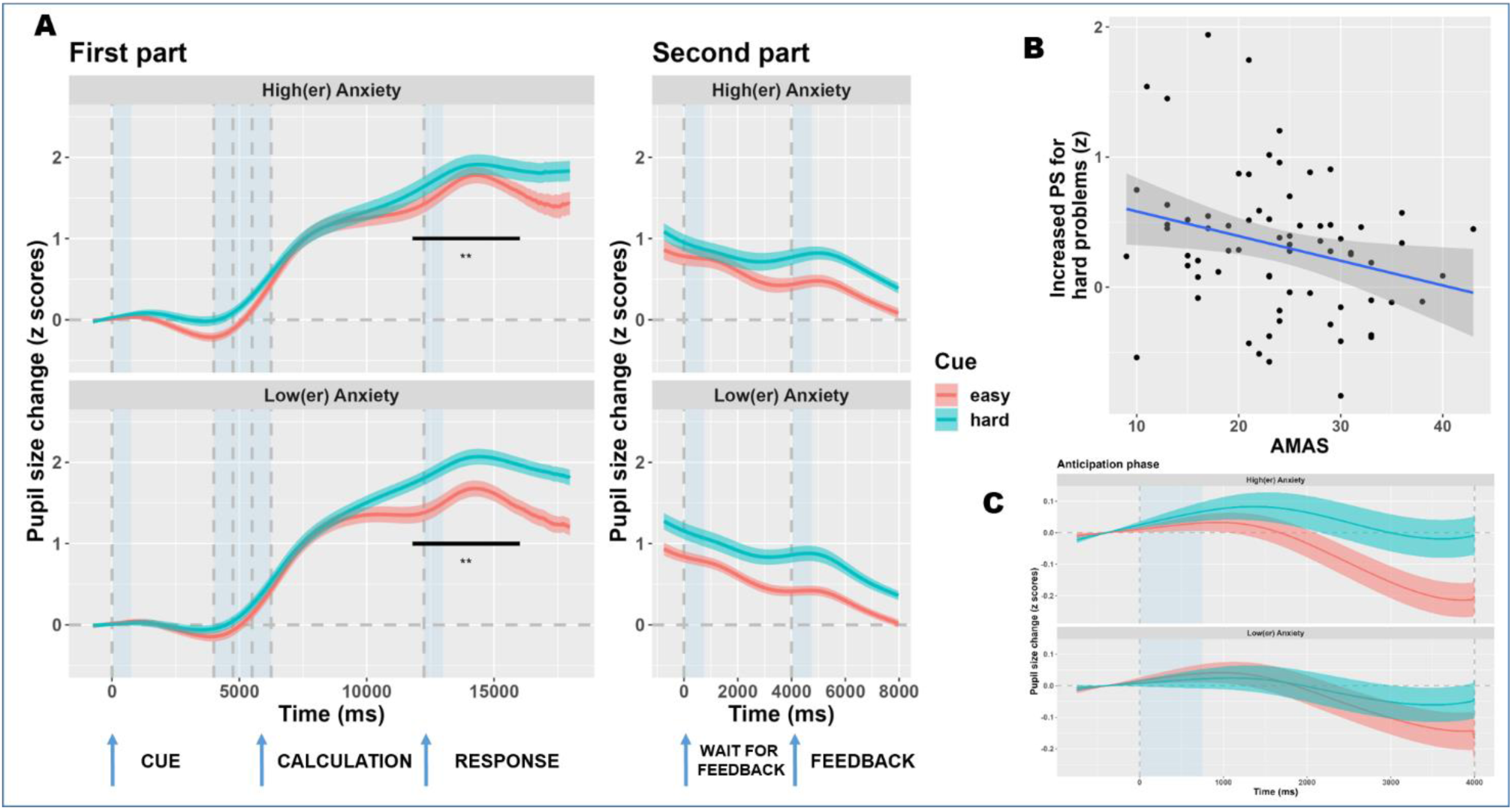
pupil size changes as a function of math anxiety. **A)** Time course of pupil size changes as a function of Math Anxiety (median split of the entire sample for depiction purposes). Results show a clear effect of task difficulty in that hard problems were associated with increased pupil dilation. A smaller, but still reliable anticipation effect is also seen across the entire sample, consisting in increased preparatory dilation following the cue for hard problems. PS was larger, for easy problems, in people with high MA during the late stages of the calculations and up until a response; however, this effect did not hold when task accuracy was accounted for in the models. The lines depict the mean PS, whereas error bars depict standard errors of the mean. Dashed gray lines delineate the onset of the different experimental phases (i.e., anticipation, calculation, response, wait for the feedback, and feedback phases). Areas shaded with transparent, light blue background indicate the presentation of sounds, as per the behavioral paradigm. **B)** Details and depiction of the correlation between MA (AMAS questionnaire) and the increase in PS observed for hard trials. Positive values indicate larger PS for hard trials, negative values larger PS for easy trials, limited to a cluster identified between 11.8 and 16s. The difference is larger for people with lower MA, presumably because of the shorter duration of their cognitive efforts especially with easy trials, after which the pupils can constrict back toward the baseline. **C)** Enlarged view of the results of the anticipation phase. There was a visual trend for an increased preparatory effect for individuals with high MA, but this trend was not entirely consistent and did not reach significance.

We computed crossvalidated LMEMs using Pupil Size (PS) as dependent variable, with MA (AMAS questionnaire) and Cue (easy vs hard) as fixed factors, as well as their interaction. There was a significant main effect of Cue (*t*_(4293.88)_= 11.2, *p*<.001; peaks around 17.5 s since the trial’s onset). A clear consensus for this effect was found starting from 3.5s and lasting until the end of the trial. In particular, “hard” problems were associated with larger PS throughout the trial, starting even prior to the presentation of the numbers, which is indicative of increased alertness for this class of problems (note that participants were cued with the word “difficult”). There was no significant main effect of MA (*t*_(107.9)_= 1.55, *p*=.125). On the other hand, there was a significant interaction Cue by MA (*t*_(4663.43)_= -4.1, *p*<.001; peaks around 15.6 s since the trial’s onset, hence in the midst of the calculation/response phase). Across folds, there was a consensus for timepoints between 11.8 to 16s, and a LMEM for this window confirmed the presence of an interaction (*t*_(4688)_= -3.17, *p*=.0015). The interaction consisted in increased PS, in individuals with high MA, in “easy” trials only, such that the difference with respect to “hard” problems decreases. Indeed, there was a negative correlation between MA (AMAS questionnaire) and the average difference in pupil dilation between easy vs hard cue conditions (r_(68)_= -0.27, *p=* .024, c.i.[0.04 – 0.47], **Figure 3B**).

When math accuracy was added as additional factor in LMEMs, however, the interaction did not hold (*t*_(4388.94)_= -1.13, *p*=.26). Instead, there was a significant effect of Accuracy (*t*_(4749.32)_= -6.74, *p*<.001; peaks in the second part, feedback phase) with a consensus between 14.2s and until the end of the trial. Errors were associated with increased PS with respect to correct answers. This effect was not modulated by MA (*t*_(4691.17)_= -1.22, *p*=.22). Furthermore, there was no three-way interaction between Cue, MA, and Accuracy (*t*_(4716.61)_= 1.41, *p*=.16). To summarize, task performance (i.e., accuracy) have a significant impact on pupil size, and, when this is accounted for, the specific role of math anxiety appears negligible. We repeated the analyses above with a much more liberal approach (i.e., LMEMs for each time point and without any correction for multiple comparisons). When Accuracy was included as fixed effect, math anxiety did not modulate PS neither alone (minimum p= .146) nor in interaction with the Cue condition (minimum p= .07).

We then focused more extensively on the anticipation phase (**Figure 3C**) because of its theoretical relevance but also because differences in this phase, in principle, cannot be explained by cognitive effort, in that this phase occurs well before the presentation of numerical stimuli and thus the onset of mental calculation. Despite a visual trend suggesting that people with high MA may present an increased alertness effect (i.e., larger PS differences between hard and easy cues), this interaction did not reach significance when using the most liberal approach (i.e., intercept-only LMEMs for each time point, not accounting for task accuracy: minimum p= 0.2).

The results outlined above only apply to Math Anxiety. Test Anxiety (TAI questionnaire: *t*_(4689.13)_= - 0.6, *p*=.55) or Trait Anxiety (STAI questionnaire: *t*_(4614.66)_= 0.78, *p*=.44) did not interact with the Cue condition, even when Accuracy was not included in the models, suggesting a specific role of MA (although the latter was completely hidden by accuracy rate when this was among the predictors).

### Can pupil size inform about math anxiety?

In the previous section we found that math anxiety (but not test or trait anxiety) modulates pupil size; however, this modulation can entirely be explained by arithmetic performance, in that when accuracy was added in the models the modulation did not hold anymore. Here, conversely, we sought to assess whether information about pupil size can nevertheless explain part of the variance of the anxiety questionnaires, beyond task accuracy. We tested the role of five predictors: task performance (accuracy) and mathematical competence (PMP); baseline pupil diameter, maximum pupil dilation, and latency of the peak (pupillary indices). In all analyses, the BIC for the null model (intercept-only) was BIC= 206.14 (note that anxiety scores were z-transformed prior to modelling, and thus the BIC for the null model remains the same across all questionnaires). Results are reported in **Table 2**.

**Table 2:** contribution of mathematical performance and pupil’s peak latency in explaining anxiety scores. The table reports the main statistics of the models compared. Task accuracy was indeed a good predictor of math anxiety, explaining a significant part of the variance of the AMAS questionnaire, and consistently with the fact that math anxiety and math performance are strictly intertwined. The latency of the pupillary peak response, indexing a more prolonged effort, was however capable to explain additional variance, up to 23% in left-out individuals. The effect of mathematical performance was only seen for math anxiety, confirming the specificity of this domain. On the other hand, a more prolonged effort was also seen in individuals with higher test anxiety scores, albeit less pronounced than that observed in the specific domain of mathematics. None of the variables were associated, in our sample, to general, trait anxiety, suggesting that prolonged cognitive effort may be a signature of the more performative situations, and that in these cases anxiety may hamper an efficient processing of information.

| **Model** | **AMAS:**<br>Accuracy | **AMAS:**<br>Accuracy +<br>Latency | **TAI:**<br>Accuracy | **TAI:**<br>Accuracy +<br>Latency | **STAI:**<br>Accuracy | **STAI:**<br>Accuracy +<br>Latency |
| --- | --- | --- | --- | --- | --- | --- |
| **Adj-R2** | .12<br>[.01-.30] | .28<br>[.14-.48] | .04<br>[0 -.17] | .17<br>[.04-.38] | 0<br>[0 -.06] | 0<br>[0 -.14] |
| **cv-R2** | 0.08 | 0.23 | 0 | 0.13 | 0 | 0 |
| **BIC** | 200.72 | 189.33 | 206.72 | 199.25 | 210.25 | 213.26 |
| **BF (vs**<br>**null)** | 15.04 | 4469 | 0.75 | 31.35 | 0.13 | 0.22 |

For math anxiety, the model only including task accuracy presented a BIC of 200.72, far superior to the null model (BF= 15). Accuracy alone accounted for 11.6% of the variance in MA scores, with a predictive performance reaching 8.4% of the variance (of left-out individuals). The best model, however, was achieved when the latency of peak PS was included among the predictors, and included both latency and task accuracy (BIC= 189.33). This model was far better than the null model (BF= 4469) but also better than the model only including task accuracy (BF= 297.4); the model including both was favored with respect to the one only including latency (BIC= 191.71, BF= 3.27). This final model consisting of two predictors reported indeed a much better performance (F_(2, 67)_=14.59, *p<*.001, adjusted-R^2^= 0.28), which translated to improved predictive power (cv-R^2^= 0.23). In summary, not much the extent by which the pupils dilate, but rather the duration of cognitive effort was a good predictor of MA, even beyond the reported task accuracy. This is in agreement with the observed correlation between this value and MA as outlined in the paragraph above (**Figure 2C**).

We repeated the analyses above for Test Anxiety (TAI questionnaire). The BSR approach found, in this case, that the best model only included peak latency among the predictors (F_(1, 68)_=14.69, *p<*.001, adjusted-R^2^= 0.17, BIC= 196.7); this model was favored by a factor BF= 112.3 with respect to the null, intercept-only model. Differently from math anxiety, performance (accuracy) was not a good predictor of test anxiety in our sample and paradigm (BIC= 206.72, BF= 0.75, thus favoring the null model). The model including both factors achieved a good performance (BIC= 199.25, BF= 31.36, cv-R^2^= 0.13), though it was 3.6 times less likely than the model only including latency. This latter model is also described in **Table 2** for comparison with the results obtained for math anxiety. In summary, unlike math anxiety, the behavioral performance (task accuracy) was not as a good predictor of test anxiety scores. The duration of cognitive effort, on the other hand, as indexed by the latency of pupillary peak dilation, also had a role in explaining part of the variance in the TAI questionnaire.

Finally, when we attempted to predict the scores of the STAI (trait anxiety) questionnaire, no model was better than the null model (BIC= 206.14). The second best model included only one predictor, peak latency, and it was not superior to the null model (BIC= 209.07, BF= 0.24, suggesting that the null model was 4.24 times more likely). Indeed, the model was not significant (F_(1, 68)_=1.33, *p=*.25) and the performance was very poor (adj-R^2^= 0.004, cv-R^2^= 0). Thus, the latency of peak PS did not add significant information apt to better identify people with higher trait anxiety, unlike the more performative domains such as test or math anxiety.

## Discussion

In this study, we measured Pupil Size (PS) in young adults with various degree of anxiety while engaged in mental calculation, as well as during the anticipatory stage and during the feedback. We have found that individuals with high levels of Math Anxiety (MA) presented, on average, increased PS, especially for easier calculations, starting a few seconds after the presentation of the second number, thus well after mental calculation started. This finding was specific for Math Anxiety (MA), and did not translate to test anxiety or trait anxiety. It is therefore very tempting to ascribe these results to the fact that MA may interfere with the processing of symbolic arithmetic, just as postulated by important models (Ashcraft & Kirk, 2001). MA may create a dual-task setting, hampering the efficient processing of information; this effect may be more pronounced with easy(er) calculations because, unlike hard(er) problems, PS generally does not reach its maximum size (i.e., ceiling), thus leaving room to appreciate a modulation. The picture is, however, much more nuanced, and while an increased cognitive effort due to MA cannot be entirely dismissed, mathematical performance was clearly the major modulating factor. In line with classic studies (Ahern & Beatty, 1979), we indeed found both larger PS and delayed PS peaks in individuals with low task performance, which generally presented much higher MA levels.

There was a substantial overlap between MA and math accuracy, so that students with high MA generally performed worse. Previous studies have found that children with developmental dyscalculia can be twice as likely to have high mathematics anxiety; however, almost 80% of children with high mathematics anxiety have typical or high mathematics performance (Devine et al., 2018). Our study does not question the existence of this dissociation. In fact, almost all participants performed quite well here. Still, the association was relevant enough that an unsupervised clustering approach found a remarkable separation between these two dimensions: participants were grouped together based on their having both high MA *and* lower accuracy (or viceversa). This suggests that assuming a substantial correlation between MA and performance (as in our own data) is rather parsimonious, on the one hand, and that attempting to match participants for their mathematical performance would probably be artificial and lacking ecological validity, on the other hand. In other words, MA and arithmetic competence would be, in young adults, too rigidly intertwined to be easily discernible.

This connection is likely due to an underlying, upstream connection between MA and numeracy, or overall mathematical competence. Indeed, our study confirmed the strict link between MA and numeracy, in that people with high MA generally presented lower scores in the psychometric battery evaluating several pillars of mathematical reasoning, beyond mental calculation. It is very difficult to establish a causal direction between these two factors (Carey et al., 2016): on one hand, MA may create a significant barrier for the development of adequate mathematical competences; on the other hand, less developed mathematical competences, through repeated failures, may ultimately lead to high MA levels. The two accounts are not mutually exclusive, so that the cause-consequence debate remains difficult to solve in absence of longitudinal and interventional studies (Carey et al., 2016). At any rate, our findings corroborate and stress the importance of studying the interaction between MA and mathematical competences early on: tackling this connection before it is too rigidly crystalized may be achieved through early educational interventions.

In a group of young adults, it is instead challenging to rule out the specific effect of MA from that of core individual differences related to the efficiency by which information is manipulated. The finding of increased PS, in people with high MA, can indeed be largely explained by the duration of the required cognitive effort. People with high MA (and lower efficiency) generally need more time to reach a solution, thus sustaining pupillary dilation for longer; people with lower MA (and higher efficiency) generally reach earlier the same solution, and are therefore free to interrupt their cognitive efforts (hence, pupils can constrict back to the baseline earlier). This is an aspect about which pupil size can inform us greatly. We used the time of peak PS as a proxy measure for the duration of cognitive effort: this measure predicted well the scores obtained in the questionnaire evaluating MA, well beyond task accuracy (which was also an important predictor). We also found that, when accuracy is accounted for, the extent of pupillary dilation (i.e., the maximum observed dilation) does not appear to have a role in predicting MA. This seems to suggest that MA may be more tightly linked – rather than to a different extent of cognitive effort and cognitive load – to a slower, less efficient information gathering and processing. Peak latency was also a good predictor of test anxiety, but not of trait anxiety; this link is therefore not specific for the mathematical domain, but rather extend to more performative situations. There are several ways in which anxiety can be distracting, all while not increasing the overall amount of cognitive effort. One possibility is that, under pressure and test situations, attentional resources (broadly defined) may be diverted away sufficiently to hamper an optimal flow of cognitive processes, thus causing mental efforts to be prolonged in time rather than accentuated. All in all, this can be taken as a further warning about the long-term impact of anxiety, in both the mathematical and broad performance domain.

For what concerns the additional question of our study – i.e., whether pupil size can be leveraged to complement and possibly improve our identification of MA – results are also mixed. On one hand, the latency of pupillary responses could significantly improve the prediction of MA scores, beyond the mere mathematical accuracy and competence. On the other hand, it is unclear what the added value of this measure would be with respect to a simpler measure of response times, which may also represent a (more convenient) proxy for processing speed. Furthermore, this signature may not be as specific, being associated more broadly to test and performance anxiety, and thus fail to differentiate worries that are specific to mathematics. Math and test anxiety are, however, positively correlated, and thus this latter finding is to be expected regardless. All in all, thus, pupil size cannot yet be regarded, alone, as a useful biomarker of MA, at least in young adults. However, more studies are warranted to explore its potential as a tool to evaluate MA in the context of different paradigms or in children. Here, pupillometry may be more sensitive in highlighting the early emergence of associations between MA and mathematical performance, through some initial failures, before these have become too entrenched to be discernible. In order to do so, beside acquiring a reliable, established physiological signal, such as PS, we stress the importance of an appropriate behavioral paradigm, encompassing stages of mathematical reasoning that extend beyond mental calculation, in particular anticipatory and feedback-related processes.

## Acknowledgements

This publication was produced with the co-funding European Union - Next Generation EU, in the context of The National Recovery and Resilience Plan, Investment 1.5 Ecosystems of Innovation, Project Tuscany Health Ecosystem (THE), CUP: B83C22003920001. The authors would like to thank Margherita di Gaudio for her assistance with data collection.

## Author contributions

**Elvio Blini:** Conceptualization; Methodology; Software; Formal Analysis; Investigation; Data curation; Visualization; Writing – original draft, review and editing

**Giovanni Anobile:** Conceptualization; Methodology; Writing – review and editing; Supervision;

**Roberto Arrighi:** Conceptualization; Methodology; Writing – review and editing; Supervision; Project administration; Funding acquisition.

## Research Transparency Statements

The authors declare no conflicts of interest.

Preregistration: The hypotheses, methods, and analysis plan were not preregistered. Materials: the script to implement the eye tracking task is publicly available. Data: all primary and preprocessed data are publicly available. Analysis scripts: all analysis scripts are publicly available. Link to the repository: https://osf.io/szb24/

Ethics: the experimental procedures were approved by the local ethics committee (Commissione per l’Etica della Ricerca, University of Florence, July 7, 2020, n. 111). The research was carried in accordance with the Declaration of Helsinki, and informed consent were obtained from all participants prior to the experiment.

